# Plasticity facilitates rapid evolution

**DOI:** 10.1101/2022.05.04.490584

**Authors:** Mikhail Burtsev, Konstantin Anokhin, Patrick Bateson

## Abstract

Developmental plasticity enables organisms to cope with new environmental challenges. If deploying such plasticity is costly in terms of time or energy, the same adaptive behavior could subsequently evolve through piecemeal genomic reorganisation that replaces the requirement to acquire that adaptation by individual plasticity. Here we report a new dimension to the way in which plasticity can drive evolutionary change leading to ever greater complexity in biological organization. Our model deploys the concept of partially overlapping functional systems. We found that plasticity accelerated dramatically the evolutionary accumulation of adaptive systems in model organisms with relatively low rates of mutation. The effect of plasticity on the evolutionary growth of complexity was even greater when the number of elements needed to construct a functional system was increased. These results suggest that as the difficulty of challenges from the environment become greater, so plasticity exerts an ever more powerful role in meeting those challenges and in opening up new avenues for the subsequent evolution of complex adaptations.

## 1 Introduction

The role of plasticity in the development and evolution of both plants and animals is attracting a great deal of current interest [1–8]. Ideas about the role of plasticity in evolution have had a long history. The most famous hypothesis is thought to have originated with Baldwin [9], Lloyd Morgan [10] and Osborne [11] who independently suggested that an individual’s adaptability could prepare the ground for an evolutionary change in which the same outcome is eventually achieved without such plasticity. However, the idea was originally proposed 23 years earlier by Spalding [12]. Bateson [13] has suggested, therefore, that the misleading term “Baldwin effect” should be replaced by the descriptive term “Adaptability driver”. The mechanism for how plasticity could influence evolution has intrigued many authors [8, 14–23]. West-Eberhard [24] has been a prominent advocate of the view that an individual’s plasticity played an important role in evolution and she has been supported by other influential writers [25].

The hypothesis that adaptability drives evolution has been repeatedly modeled, both analytically [25–31] and by simulation [31–37]. Nevertheless, the hypothesis is widely supposed to be of limited interest because it merely proposes a mechanism by which one phenotype acquired through the organism’s adaptability is replaced, in the course of Darwinian evolution, by an inherited mechanism that expresses itself as a phenotypic copy at lower cost. The hypothesis was not thought to provide a general explanation for evolutionary processes [38].

The standard explanation given for the evolution of complex adaptive behaviour is that it involved the gradual accumulation of the necessary component processes by reorganization of the genome alone [39, 40]. The problem is that in most cases even the evolution of the simplest adaptive trait requires a number of independent reorganisations of the genome. When the adaptation depended on the simultaneous occurrence of a combination of such reorganisations that would be useless on their own, inevitably the evolutionary process would have been slow. Moreover, the time to acquire the adaptation by mutation grows nonlinearly with the required number of genomic reorganisations. Suppose that every reorganisation is expected to appear in a population every 10 generations then the addition of one more reorganisation to the combination will delay appearance of the adaptation 10 fold. Yet a growing body of evidence suggests that complex behaviour has evolved rapidly in birds and mammals [41–44]. In this paper we show how such integrated systems could appear rapidly in the course of evolution under the guidance of an individual’s plasticity and the use of elements shared by different functional systems. Many different forms of plasticity have been recognized, including the various forms of learning operating at the behavioural level, while other forms of plasticity operate at lower levels of organisation [3]. We argue that the effect of plasticity on evolution became increasingly powerful as animals became more complex. As components of functional systems requiring plasticity are genetically assimilated in the course of evolution then less and less plasticity is required to integrate inherited elements into the functional system. If the capacity for plastic change remains constant, it can be used by an animal to acquire other, previously unavailable, adaptations. This “assimilate-stretch” process [22] creates constant pressure for further assimilation and for retaining plasticity. In our study we examine how such a process guided by plasticity and relying on partial overlap between different functional systems leads to the accumulation of complexity in biological organisation.

## 2 The model

We have incorporated into our simulation the concept of “functional systems” [45]. The general idea is that the fitness of an individual depends on systems organized as different combinations of phenotypic elements such as connections in the neural network, the capacity to use information contained in the energy impinging on a sense organ, specific biochemical reactions, and particular effectors that respond adaptively to the stimulation. Elements may be recombined in different ways to perform different functions. Novel challenges create the conditions for the emergence of new functional systems added to the existing ones either by Darwinian evolution or by an individual’s plasticity. Possible examples are the addition of a face recognition module in primates [46] and the evolution of habitat invasiveness in birds [42]. This evolutionary process leads to the establishment of an increasingly elaborate phenotypic organization and patterns of behaviour. When such complexity entails a greater ability to discriminate between different features of the environment or a greater ability to manipulate the environment, the organism will benefit and will be more likely to survive and reproduce in the face of multiple challenges during its lifetime.

A new inherited adaptation emerges in evolution when the accumulated phenotypic effects of genomic reorganisation are added to the existing phenotype. Although these phenotypic effects are specific to the new function, existing parts of the phenotype are also recruited for this function. As a result, phenotypic elements established earlier in evolution should be incorporated into more adaptive systems than later evolved elements. To simulate this process, functional systems in our model are nested in a hierarchy from old to new.

The model organism in our simulations is made up of functional systems and each system consists of a number of elements (Fig. 1). Definitions of all the parameters and variables used in the model are given in Table 1. The phenotype is described by the set 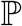 of *f* binary strings given by **e**^*i*^ for the *i*th functional system:

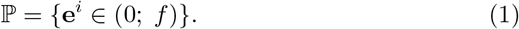

**Fig. 1.**
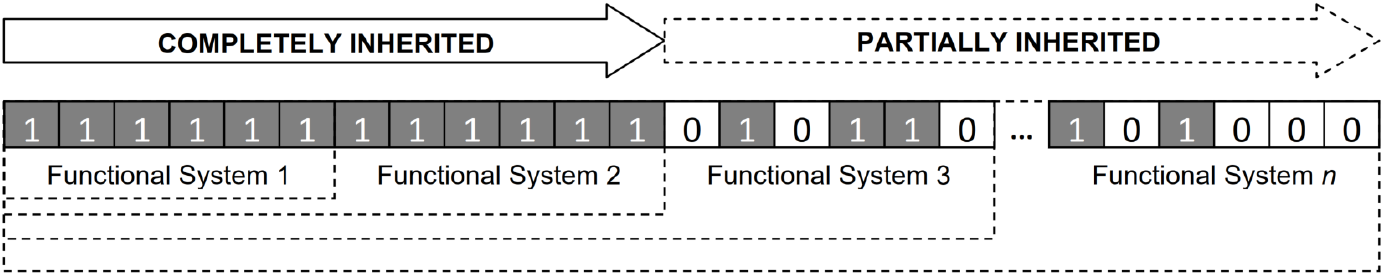
The model of an organism capable of learning. Each functional system consists of elements, the number of which is specified by a parameter (*C*). An element is either present (‘1’) or absent (‘0’). Each inherited element is influenced by a gene. An absent element may be made operational (‘0’ converted to ‘1’) by genetic mutation or by learning. In each individual model organism development proceeds step by step through fully inherited systems until a system is reached in which an element is missing. In members of populations that can learn an element may be made operational by a learning process that replaces a ‘0’ with a ‘1’ over fixed number of trials specified by a parameter (*t*). The organisms with the largest number of complete functional systems are most likely to survive and reproduce.

**Table 1.** Definition of parameters and variables used in the model.

| Parameter | Description | Value(s) |
| --- | --- | --- |
| $N$ | population size | 500 |
| $n_g$ | number of generations in simulated evolution | 25000 |
| $L$ | number of loci in genotype | 4000 |
| $A$ | number of alleles per gene locus | 5 |
| $\mu$ | probability of point mutation | $10^{-6} - 10^{-3}$ |
| $f$ | number of functional systems per individual in the simulation | 2000 |
| $C$ | number of phenotypic elements required to acquire a new functional system (i.e. the complexity of new system acquisition) | 10, 20, 40 |
| $p_e$ | probability of filling in missing phenotypic elements in the terminal system during learning trial | 0.5 |
| $t$ | number of learning trials | 0, 1, 5, 10 |
| $\mathbf{e}^i$ | a string of zeroes and ones that describes the composition of the $i$th functional system | |
| $m$ | he number of missed elements in an incomplete functional system | |

Here *f* is the maximal possible number of functional systems per organism during the simulation. The length of the string is given by the parameter *C*. Components of the given binary string encode the presence of the given feature in the phenotype. If the value of a component element is 1, then the corresponding feature is included in the phenotype; otherwise it is not. A new system is composed by the addition of *C* elements to the terminal system; thus the functional system *FS_j_* includes all strings **e**^*i*^ with *i* ≤ *j*:

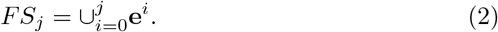

Where 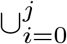 is a union operator representing the concatenation of binary strings with indexes from 0 to *j* to form representation of the *j^th^* functional system in the model. Each element is necessary for the working of the whole functional system and the system will not become functional until it has a complete set of elements, i.e. all their bits are set to 1. Only completed systems contribute to the organism’s fitness. The genetic architecture of each individual specifies the states of each of its systems’ elements. The genotype of the individual consists of *L* loci with *A* alleles per locus. Every allele is connected to some phenotypic element and if this allele is expressed in the genotype then the corresponding element is manifested in the phenotype.

The fitness of the phenotype is equal to the number of complete systems. In the real world, the complexity of an organism can decrease in the course of evolution, as in the case of many parasites, but in the cases we have sought to model, the capacity to handle a large number of different challenges by the environment carries a distinct advantage.

The parent’s genome is inherited by the offspring, so their phenotypes are the same except that each element can be changed by a point mutation in a gene relating to that element. Genomic reorganisation can add or remove an element from a functional system. For simplicity we have represented such reorganisation as a single point mutation.

An individual in the model is initialized by the sequential assembling of systems from inherited elements that terminates when the first system with a missing element is encountered. If due to mutation some phenotypic element of an intermediate system is lost, then the developmental progression is broken at this stage and none of the functional systems at subsequent developmental stages are able to make contributions to fitness. On the other hand, a mutation can produce an element that completes a new terminal system and increases the organism’s fitness. The difficulty of acquiring a new system is affected by the parameter *C* that determines the number of elements required for a complete system.

A new system can be also generated within an individual in those populations in which plasticity is possible. Plasticity can switch elements from 0 to 1. During the period of plasticity an individual is allowed to perform one attempt to fill in with probability *p_e_* a missing phenotypic element in the terminal system. Hence, the more elements that are missing, the less likely is plasticity to be successful. A new system can be also generated within an individual in those populations in which plasticity is possible. Plasticity can switch elements from 0 to 1. During the period of plasticity an individual is allowed to perform a fixed number of attempts to fill in independently with probability *p_e_* missing phenotypic elements in the terminal system. Given *t* trials an individual is able to complete one system per trial with probability:

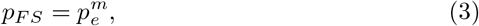

where *m* is the number of missed elements in the system *FS*. Hence, the more elements that are missing, the less likely is plasticity to be successful. If the trial was successful, remaining trials are utilized for further development by probabilistic addition of subsequent functional systems. Acquired elements are not inherited; Lamarckian inheritance does not occur in this model. However, later in evolution, the element filled by plasticity may be replaced by genomic reorganisation, simulating the adaptability driver. In our model the replacement by genomic reorganisation of an element previously filled by plasticity releases the descendents of that organism to meet other challenges set by the environment.

When the phenotype of each individual in the population has been specified, *N* offspring that will constitute the next generation are obtained. A potential candidate for breeding is selected at random from the population. The probability *p_rep_* that it will reproduce is equal to its number of completed functional systems relative to the individual in the population with the largest number of functional systems:

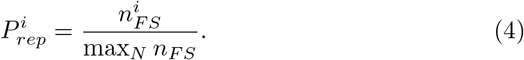

Here 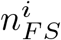 is a number of completed systems for *i^th^* individual; *N* is a population size. The offspring has the same genotype as the parent before point mutation is applied to each locus with probability *μ*. A summary of the simulation routine is given by Algorithm 1. We studied two variants of the model with and without learning. In particular we explored how the rate of phenotypic evolution in the both variants depends on mutation rate, the difficulty of new system acquisition and the number of learning trials.

### Algorithm 1 Summary of the simulation procedure.

1. The genotype-phenotype map is created by random assigning of alleles to phenotypic elements.
2. The population of *N* individuals is initiated. Values of loci of an individual’s genotypes are set randomly.
3. For every individual perform:
  a. Development of inherited systems:
    i. Reset all bits of the phenotype binary string to 0.
    ii. Make the bits of the phenotype string correspond to the alleles found in the genotype.
    iii. Find completed functional system with largest number of elements.
  b. Learning (in populations capable of learning):
    i. With *i* systems complete conduct t trials to complete the system *i*+1 with probability population select a parent with probability proportional to the number of its complete systems after learning.
    ii. Copy the parental genotype to the offspring and apply point mutations with probability μ per locus
4. Repeat step 3 for *n_g_* generations.

## 3 Results

We first studied the dynamics of evolution of model organisms in populations with and without plasticity. Figure 2a shows how, for a fixed and relatively low mutation rate, the capacity to learn greatly increases the number of sequentially ordered systems that evolve over 25,000 generations. The rate of acquisition of systems starts to level off because, in this model, the greater the number of systems that have been accumulated, the more likely will the developmental process be disrupted by mutation. The overall result is robust across a wide range of parameter values (see below). Populations with *C* = 20 and without plasticity gained on average 14.3 systems by the end of 10 runs of simulation with mutation rate per gene locus *μ* = 0.0002. The simulations with plasticity finished with the average of 45.1 systems for the same parameter settings, a rate that is more than three times greater than in the non-plastic population. Therefore, in this model plasticity promotes much more rapid evolution of complex sets of adaptive systems than can be accomplished by mutation alone. This occurs as previously plastic elements are replaced by inherited elements and the model organism is able to fill by plasticity missing elements in subsequent systems (see Fig. 1). The proportion of inherited elements in systems beyond the last complete functional system is highest in the incomplete systems closest to it (Fig. 2b).

**Fig. 2.**
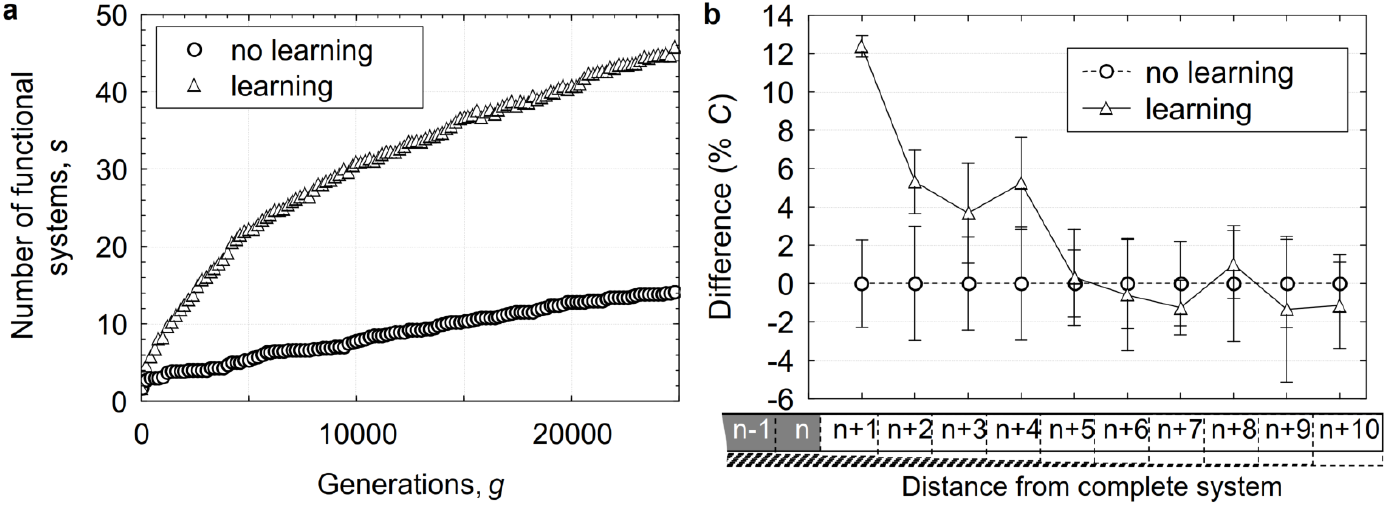
**a. Effects of plasticity on the rate of evolution.** The number of inherited functional systems acquired during evolution increases more rapidly in populations that exhibit plasticity (triangles) than in populations that do not (circles). In each generation the mean of 10 simulations is shown for the individuals with the largest number of functional systems. **b. Differences between the acquisition of the elements of functional systems in plastic and non-plastic populations.** The proportion of inherited elements in systems beyond the last complete system is significantly higher in plastic populations, as shown by 95% confidence limits, than in non-plastic populations. The difference between populations decreases with stages more remote from the last complete functional system. For results on the both panels the number of elements required to generate a functional system, *C* = 20 and the mutation rate per gene locus, *μ* = 0.0002.

The effect of plasticity on evolution of complex biological structure and behaviour in the model is even more dramatic if the number of generations to evolve the same amount of functional systems is considered. For the same settings as used in Fig. 2, it took 2458 generations for the plastic populations to accumulate 14.3 systems. This is more than ten times faster than in the populations evolving without plasticity. To investigate how the capacity of plasticity to speed up evolution is related to the complexity of the task we conducted simulations by varying the number of elements in each system. The difficulty of acquiring a new system in our model is controlled by the parameter *C*. The value of this parameter determines how many elements are required to make a new system functional. We studied the populations with *C* equal to 10, 20 or 40. The ratio of the generations required to evolve the phenotype with the same number of functional systems in populations with and without plasticity is given in Fig. 3a. For both populations with and without plasticity, the rate of evolution decreased with the task complexity but on average the relative potentiating effect of plasticity was four times greater for *C* = 40 than for *C* =10 (Fig. 3a). In the case of the most complex task, plasticity accelerated accumulation of adaptive systems about 40-fold. On the assumption that the number of functional systems is related to the complexity of biological organization, these results suggest that plasticity accelerates the evolution of yet more complex organisation. In the model the acceleration is greater as more elements have to be added to the systems and the difficulty of completing a terminal system is increased. The chances of members of nonplastic populations acquiring solutions to the most complex of problems are much lower than in members of plastic populations, and if they did acquire them, would require a very long period of evolution.

**Fig. 3.**
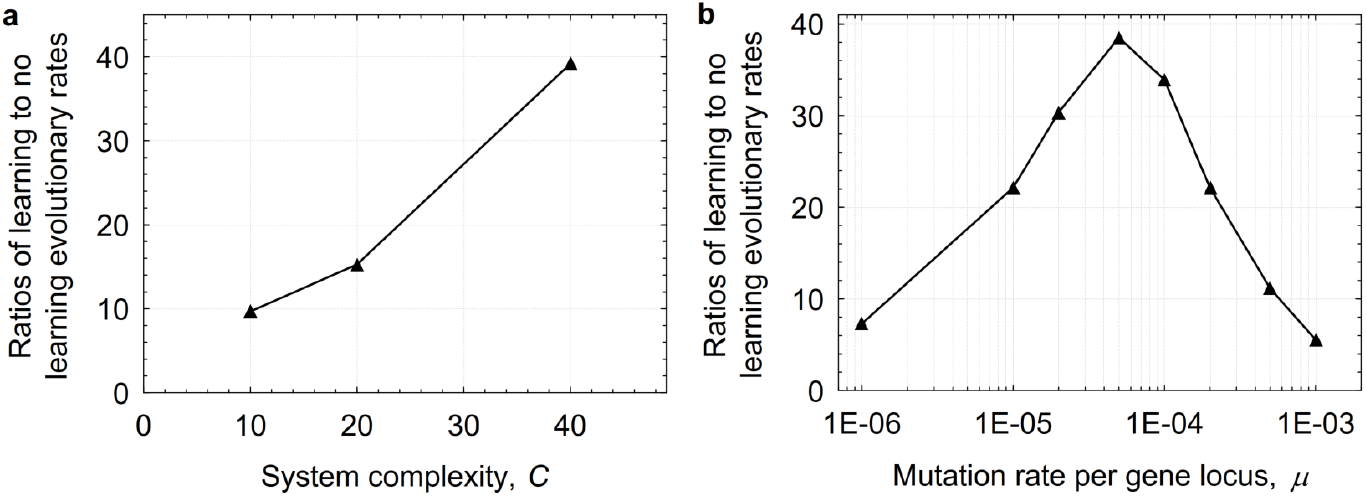
Effects of varying system complexity on rates of evolution in plastic and non-plastic populations. **a.** The acquisition rate in populations that exhibit plasticity relative to those that do not is shown for different values of the parameter C that specifies the number of elements required to generate the functional system and provides a measure of the system’s complexity. The ratios are obtained from the mean of 10 simulations and averaged over the range of the system’s complexities (*C* = 10, 20, 40). **b.** The acquisition rate in populations that exhibit plasticity relative to those that do not is shown for different values of the parameter, μ, for different rates of mutation. The ratios are obtained from the mean of 10 simulations and averaged over the range of mutation rates (*μ* = 10^−6^ − 10^−3^).

Next we studied how the effect of adaptability on evolution depends on the rate of mutation (see Fig. 3b). With increasing mutation rate from *μ* = 10^−6^ to 10^−3^, the acceleration of evolution by plasticity first grows to a maximum at *μ* = 10^−4^ and then decreases. For high mutation rates, acceleration decreases due to disruption of already established adaptive elements. For lower rates of mutation the effect of mutation rate on the difference between the plastic and non-plastic populations is explained by the likelihood that a plastic element will be replaced by an inherited one.

Finally we studied how the rate of evolution changed with increasing number of learning trials in populations capable of learning (Fig. 4). Even one learning trial is sufficient to accelerate significantly the accumulation of adaptive phenotypic modifications. Populations with 2 and 5 learning trials evolve faster than populations with 1 trial.

**Fig. 4.**
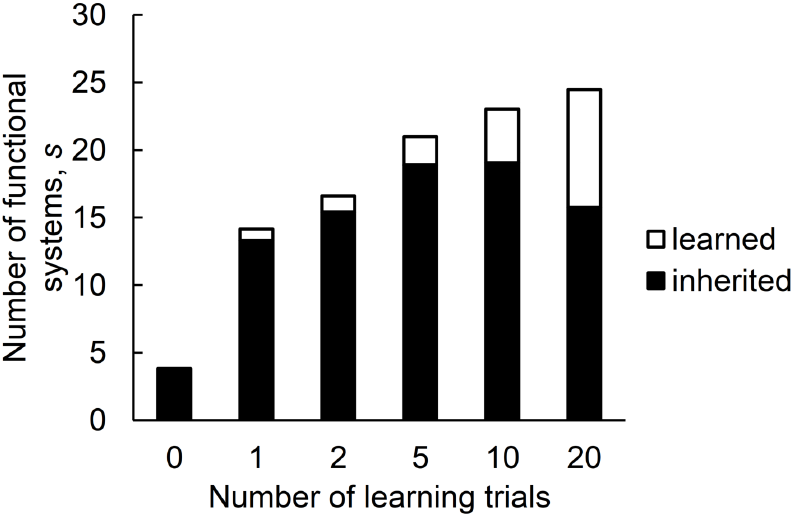
Effect of amount of learning on evolution. The number of functional systems after 25,000 generations with varying amount of learning specified by the number of learning trials *t*. The data are obtained from the mean of 10 simulations and averaged over the range of mutation rates (*μ* = 10^−5^ − 10^−3^) with the complexity *C* = 40.

## 4 Discussion

In as much as it has been taken seriously, the adaptability driver has usually been taken as providing a mechanism for the slow accretion of spontaneously expressed phenotypic elements in the course of evolution [38]. Emphasis was placed on how particular behaviour patterns initially acquired by learning could be expressed spontaneously without learning in the course of subsequent evolution [10]. Recent developments initiated by the work of Hinton and Nowlan [32] have shifted the focus to other issues, such as, the way in which plasticity can accelerate the rate at which challenges set by the environment can be met [27, 28, 31, 47, 48], the advantages of plasticity in a changing environment [29, 33, 36, 48–52], and the conditions in which plasticity might slow down evolution [27, 31]. Our study complements such proposals about the role of plasticity in evolutionary processes. However, the study adds a much more important feature to the general discussion of developmental plasticity and evolution. By incorporating Pyotr Anokhin’s [45] concept of functional systems, the simulations indicate how plasticity could have facilitated rapid evolutionary change. From the perspective of a single ecological challenge requiring just one functional system, our model is similar to a classical single-peaked landscape simulation [32]. However, the main highlight of our model is its operation in a complex ecological landscape in which many challenges face the organism [53]. It allowed us to examine the role of plasticity in the evolution of multiple partially overlapping functional systems. We have not been concerned here with how the capacity to learn from others affects genetic change which has been the central focus of Boyd, Richerson and their colleagues [54].

Evolutionary theory has provided elegant explanations for ways in which complexity might be gradually elaborated. The many steps from simple light detectors to complex vertebrate eyes have been well described [55, 56]. Major transitions in evolution have been explained in terms of changes in genetic regulation early in development [57] and these explanations have been offered for the explosion of variety seen in the Cambrian [58]. Our study suggests another way in which the process of rapid evolution might have been driven, particularly in more complex animals. The model demonstrates how mutations each of which produces small variation can be accumulated under the guidance of plasticity to create substantial adaptive change of the phenotype. The model explains how rapid evolutionary change can be squared with maintaining the overall functionality of the organism [59].

If we are correct, the role of plasticity would have become more and more important as phenotypes increased in complexity. Furthermore, if plasticity had not been possible until a certain level of complexity had evolved, then a sharp increase would occur in the rate of evolution at that point. By varying the amount of plasticity in the model we found that it affects speed of evolution in non-linear manner. Some theorists have argued that plasticity could dampen down the rate of evolution (e.g. [60, 61]). Their proposal was that, with every individual in the population coping plastically with an environmental challenge, natural selection would have had no variation on which to act. In some cases this might well have been true in the short run. However, if operating plastic mechanisms involved time and energy costs, then individuals that expressed the adaptation spontaneously would readily invade the population and the dampening effect of plasticity on evolutionary rate would be lost.

## 5 Conclusion

In general, our simulations suggest that an ability to cope with complex environmental challenges by adding to the elements of existing functional systems, plasticity opens up ecological niches previously unavailable to the organism. These plastic changes could include the creative effects of play [62]. The beneficial effects would inevitably lead to the subsequent evolution of morphological, physiological and biochemical adaptations to those previously unoccupied niches. Where an environmental challenge involved greater processing capacity by the brain, this organ too would be expected to evolve with greater rapidity. On the assumption that the bigger brain ensures greater learning capacity, the rate of evolution should correlate positively with the relative brain size. This expectation is given some support by the study suggesting that taxonomic groups evolving most rapidly have the biggest brains relative to body size [41]. The expectation is also supported by the correlation between behavioural innovation and brain size reported for birds [42] and primates [63]. As the beneficial consequences of completed functional systems became available to it, the animal would have then been able to acquire by learning adaptations to new challenges set by the environment. We conclude, therefore, that each individual’s plasticity provided an evolutionary ratchet and this provided direction towards ever greater complexity.

## Acknowledgments

The publication of this article is dedicated to the memory of Patrick Bateson. Most of the work was done in 2007-2008 when the KA and MB visited Cambridge to work with PB on a grant from the Royal Society. PB, KA and MB developed conceptual foundations for the model. MB implemented and performed computer simulations. Simulation results and additional experiments were discussed by PB, KA and MB. PB, KA and MB wrote the paper. We also want to thank Eva Jablonka for fruitful discussions of our work from its very beginning. This publication closely follows the final draft discussed with PB in January 2014, unfortunately the work was never submitted to publication after that.

For us Pat was not only an academic peer but also a friend. His passion for science and for hard questions of animal biology not only guided this work but also gave us a much deeper view of foundational problems on the intersection of evolution, development, behavior and learning.

